# Emerged human-like facial expression representation in a deep convolutional neural network

**DOI:** 10.1101/2021.05.08.443217

**Authors:** Liqin Zhou, Ming Meng, Ke Zhou

## Abstract

Face identity and expression play critical roles in social communication. Recent research found that the deep convolutional neural networks (DCNNs) trained to recognize facial identities spontaneously learn features that support facial expression recognition, and vice versa, suggesting an integrated representation of facial identity and expression. In the present study, we found that the expression-selective units spontaneously emerged in a VGG-Face trained for facial identity recognition and tuned to distinct basic expressions. Importantly, they exhibited typical hallmarks of human expression perception, i.e., the facial expression confusion effect and categorical perception effect. We then investigated whether the emergence of expression-selective units is attributed to either face-specific experience or domain-general processing, by carrying out the same analysis on a VGG-16 trained for object classification and an untrained VGG-Face without any visual experience, both of them having the identical architecture with the pretrained VGG-Face. Although Similar expression-selective units were found in both DCNNs, they did not exhibit reliable human-like characteristics of facial expression perception. Taken together, our computational findings revealed the necessity of domain-specific visual experience of face identity for the development of facial expression perception, highlighting the contribution of nurture to form human-like facial expression perception. Beyond the weak equivalence between human and DCNNS at the input-output behavior, emerging simulated algorithms between models and humans could be established through domain-specific experience.

## Introduction

Facial identity and expression play important roles in daily life and social communication. When interacting with others, we can not only easily recognize who they are through their facial identity information, but also access their emotions from their facial expressions. An influential early model proposed that face identity and expression were processed separately via parallel pathways [1,2]. Configural information for encoding face identity and expression was different from each other [3]. Findings from several neuropsychological studies supported this view. Patients with impaired facial expression recognition still retained the ability to recognize famous faces [4,5], whereas patients with prosopagnosia (an inability to recognize the identity of others from their faces) could still recognize facial expressions [4–6]. Haxby et al. (2000) further proposed a distributed neural system for face perception, which emphasized a distinction between the representation of invariant aspects (e.g., identity) and changeable aspects (e.g., expression) of faces. According to this model, in the core system, lateral inferior occipitotemporal cortex (i.e., fusiform face area, FFA; occipital face area, OFA) and superior temporal sulcus (STS) may contribute to the recognition of facial identity and expression, respectively [8,9]. Patients with OFA/FFA damage have deficits for face identity recognition, and those with damage to the posterior STS (pSTS) suffer impairments in discriminating expression [10].

On the other hand, processing mechanisms of the human visual system for facial identity and expression recognition normally share face stimuli as inputs. That is, naturally, a face may contain both identity and expression information. Early visual processing of the same face stimuli would be the same for both identity and expression recognition, but it is unclear at what stage they may start to split. Amid increasing evidence to suggest an interdependence or interaction between face identity and expression processing [11–15], we hypothesis that any computational model that simulates human performance for facial identity and expression recognition must share common inputs for training. Moreover, if domain-specific face input is necessary to train a computational model that simulates human performance for facial identity and expression recognition, it would suggest that the split of identity and expression processing might occur at or after the domain-general visual processing stages. However, if no training or no domain-specific training of face inputs were needed for a computational model that simulates human performance, it would suggest a dissociation between identity and expression processing at domain-general stages of visual processing.

Specifically, DCNNs has achieved human-level performance in object recognition of natural images. Investigations combining DCNNs with cognitive neuroscience further discovered similar functional properties between artificial and biological systems. For instance, there is a trend of high similarity between the hierarchy of DCNNs and primate ventral visual pathways [16,17]. Research relevant to this study revealed a similarity of activation patterns between face identity-pretrained DCNNs and human FFA/OFA [18]. Thus, DCNNs could be a useful model simulating the processes of biological neural systems. More recently, several seminal studies have found that the DCNNs trained for the recognition of facial expression spontaneously developed facial identity recognition ability, and vice versa, suggesting that integrated representations of identity and expression may arise naturally within neural networks, like humans do [19,20]. However, a recent study found that face identity-selective units could spontaneously emerge in a untrained DCNN [21], which seemed to cast a strong doubt on the role of nurture in developing face perception, as well as the abovementioned speculation. Indeed, when adopting a computational approach to examine the human cognitive function, a success in classifying different expressions only suggests the weak equivalence between DCNNs and humans at the input-output behavior in Marr’s three-level framework, which does not necessarily mean that DCNNs and humans adopt the similar representational mechanisms (i.e., algorithms) to achieve a same computational goal [22]. Therefore, to explore whether a common mechanism may be shared by both artificial and biological intelligent systems, a much stronger equivalence should be tested by establishing additional relationships between models and humans, i.e., identities in algorithms between them [23].

Therefore, in the present study, we borrowed the cognitive approaches developed in human studies to explore whether the human-like facial expression recognition relied on face identity recognition, by using the VGG-Face, a typical DCNN pretrained for the face identity recognition task (hereafter referred to as pretrained VGG-Face). The pretrained VGG-Face was chosen because of its relatively simple architecture and evidence supporting its similar representations of face identity to those in the human ventral pathway [18]. The training process of VGG-Face has already determined units’ selectivity for various features to optimize the network’s face identity recognition performance. If the pretrained VGG-Face could simulate the interdependence between facial identity and expression in the human brain, then it should spontaneously generate expression-selective units. The selective units should also be able to predict the expressions of new face images. However, as mentioned above, having an ability to correctly classify different expressions does not necessarily mean human-like perception of expressions. Here, we introduced morphed expression continua to test whether these units perceived morphed expression categorically in a human-like way.

Then, in order to answer what human-like expression perception depends on, we introduced two additional DCNNs. The first one is the VGG-16, a DCNN that has an almost identical architecture with the pretrained VGG-Face but was trained only for natural object classification. The other one is an untrained VGG-Face, which has identical architecture to the pretrained VGG-Face, but its weights are randomly assigned with no training (hereafter referred to as untrained VGG-Face). Comparisons among the three DCNNs would clarify whether the human-like expression perception relies on face (identity) recognition specific experience, or general object recognition experience, or merely the architecture of the network.

## Materials and methods

### Neural network models

The VGG-Face (https://www.robots.ox.ac.uk/~vgg/software/vgg_face/)[24] pretrained for recognizing 2622 face identities on a database with 2.6 million face images was used. It achieved state-of-the-art performance whilst requiring less data than other state-of-the-art models (DeepFace and FaceNet). The network consists of thirteen convolutional (conv) layers and three fully connected (FC) layers. All these sixteen layers are followed by a rectification linear unit (ReLU). The thirteen convolutional layers are distributed to five blocks. Each of the first two blocks consists of two consecutive convolutional layers followed by max pooling. Each of the latter three blocks consists of three consecutive layers followed by max pooling.

Besides, we used another two DCNNs (VGG-16 and untrained VGG-Face) for comparisons. The VGG-16 was trained for classifying 1000 object categories using the ILSVRC-2014 ImageNet database, which contains over 14 million natural visual images (https://arxiv.org/abs/1409.1556)[25]. It achieves a 92.7% top-5 test accuracy in ImageNet and thus is one of the best models submitted to the ILSVRC-2014. The architecture of the VGG-16 is identical to the pretrained VGG-Face except that the last FC layer includes 1000 units for 1000 object classes rather than 2622 units for facial identities in the pretrained VGG-Face. The untrained VGG-Face preserved the fully identical architecture of the pretrained VGG-Face while randomly assigning the connective weights (Xavier normal initialization)[18,26] without any training experience.

### Stimuli

Three stimulus sets were used in the study. Stimulus set 1 was used to detect expression-selective units in the DCNNs. It contained 624 facial expression images: 104 identities, each with 6 basic expressions (anger, disgust, fear, happiness, sadness, and surprise) [27,28]. In the stimulus set 1, seventy identities were from Karolinska Directed Emotional Faces (KDEF) database [29], and the other 34 identities from the NimStim Database [30]. Then, to validate the reliability of expression recognition capability of the expression-selective units, a second stimulus set — the Radboud Faces Database (RaFD) [31] was applied. The front view images of all the 67 identities in the RaFD were used, each identity contained the aforementioned six expressions. Furthermore, we applied AffectNet Database as the third stimulus set to test if the units could recognize facial expressions in real life stimuli. The AffectNet Database is by far the largest database of facial expressions in the wild, and its images were collected from the Internet [32]. Four thousand eight hundred manually annotated images from the AffectNet Database were used, and each expression included 800 images. In this dataset, the face identities across expressions are different.

For each dataset, the luminance and contrast of the images were matched by using the SHINE toolbox [33], and the face part were reserved with the background removed by using the facemorpher package (https://alyssaq.github.io/face_morpher/facemorpher.html). Then, the images were resized to 224 × 224 pixels.

The prototypic expressions used in the morphed expression discrimination task were from the stimulus set 1. All identities in the stimulus set 1 were used. Seven morph continua were tested in the present study, including happiness-anger, happiness-fear, anger-disgust, happiness-sadness, anger-fear, disgust-fear, and disgust-sad. The number of morphed levels in each morph continuum was 201. The morphing process was conducted by using the facemorpher package.

### Analysis of network units

Each DCNN was presented with the stimulus set 1 and the responses of the units in the final layer of the feature extraction network (conv5-3) were extracted to be analyzed. Like Nasr et al. (2019), a two-way non-repeated ANOVA with expression (6 facial expressions) and identity (104 identities) as factors was conducted to identify expression-selective units. The “expression-selective units” were referred to as those that exhibited a significant main effect of expression (*p* ≤ 0.01) but no significant effect of identity (*p* > 0.01). For each expression-selective unit, the responses were normalized across all images in the stimulus set 1. After that, its tuning value for each expression was calculated by taking the difference between the average response to all images within the same expression and the average response to all images in the stimulus set, and then dividing the difference by the standard deviation of the responses across all images in the stimulus set [35]:

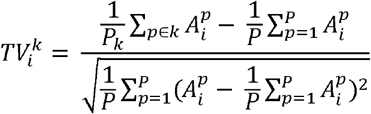

Where 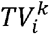 is the tuning value of unit to expression *k*. 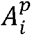 is the normalized response of unit *i* to image *p*. *P_k_* is the number of images that are labeled as expression *k*. *P* is the number of all images in the database. The tuning value reflects the extent to which a unit activates preferentially to images of a specific expression. For each unit, the expression with the highest tuning value is defined as its preferred expression.

In each DCNN, to test the reliability and generality of the expression recognition ability of expression-selective units, the SVC model trained on the stimulus set 1 was used to predict the expressions of images from the RaFD and AffectNet Database, separately. In the SVC model, the first six hundred principal components of the responses of expression-selective units were used. The prediction accuracy indicated the extent to which expression-selective units could classify facial expressions. Further, predicted expressions and true expressions of the images were used to construct the confusion matrix. And this confusion matrix was correlated with that of humans to measure how similar the expression recognition performance of expression-selective units and humans were.

### Morphed expression discrimination task

To test whether expression-selective units exhibit a human-like categorical perception of morphed facial expressions, we designed a morphed expression discrimination task that was comparable to the ABX discrimination task designed for human beings [36–38]. Taking happiness-anger continuum for example, the expression-selective units whose preferred expression was happiness or anger were selected to perform the task. At first, a binary SVC model was trained on the prototypic expressions (happy and angry expressions of all 104 identities in the stimulus set 1), then the trained SVC model was used to predict the expressions of morphed expression images (the middle 199 morph levels besides the two prototypic expressions). For each morph level, the identification frequency of anger was defined to be the network’s identification rate at the current expression morph level.

To quantitatively characterize the shape of the identification curve, we fitted Linear function, Quadratic function (poly2), and Logistic function to the curve, respectively. If the network perceived the morphed expressions like a human, the identification curve should be nonlinear and should show an abrupt category boundary. Thus, the goodness of fit (R^2^) of the Logistic function (S-shaped) to identification curves should be the best.

### Comparisons between different DCNNs

To test the dependence of human-like expression perception of expression-selective units on face identity recognition experience, we also introduced VGG-16 and untrained VGG-Face as controls. The VGG-16 is trained for natural object classification, and the untrained VGG-Face has no training experience. First, the expression classification performances of expression-selective units in different DCNNs were compared to explore whether these units in the pretrained VGG-Face recognized expressions better than the VGG-16 and untrained VGG-Face. Then, we assessed the differences of categorical perception of morphed expressions among the DCNNs by respectively comparing their goodness of fit (R^2^) of the Logistic function and goodness of fit (R^2^) of the Linear function. Finally, the Mann–Whitney U test was used for statistically evaluating the differences among the DCNNs.

## Results

### 1. Expression-selective units spontaneously emerge in the pretrained VGG-Face

We first explored whether expression-selective units could spontaneously emerge in the pretrained VGG-Face. Note that, the pretrained VGG-Face was trained for recognizing 2622 identities with over two million face images [24]. It consists of thirteen convolutional (conv) layers and three fully connected (FC) layers (Fig. 1A; details in Methods). The first thirteen convolutional layers form a feature extraction network which transforms images to a high-level representation, and the following three fully-connected layers make up a classification network that classifies images using the high-level representation [34].

**Fig. 1.**
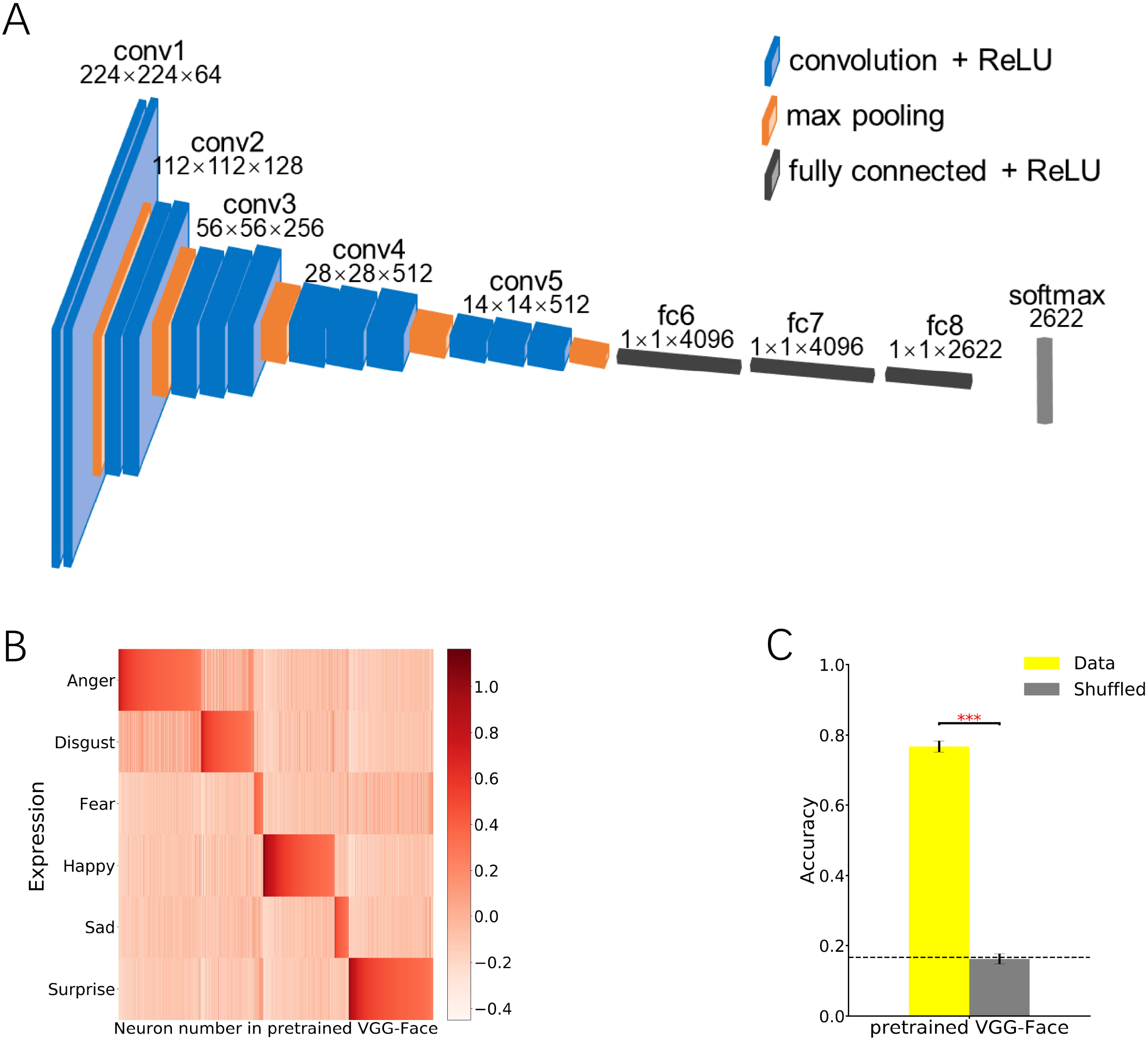
Expression-selective units emerged in the pretrained VGG-Face. (A). The architecture of the VGG-Face. (B) The tuning value map of the expression-selective units in the pretrained VGG-Face. (C) The expression classification performance of the expression-selective units. The black dashed line represents the chance level. Error bars indicate SE. ***: p≤0.001.

Here, we tested the expression selectivity of each unit in the final layer of the feature extraction network (conv5-3) with stimulus set 1 (details in Methods). The stimulus set 1 is composed of 104 different facial identities selected from the Karolinska Directed Emotional Faces (KDEF)[29] and NimStim [30] databases, and each identity includes 6 kinds of basic expressions (i.e., anger, disgust, fear, happiness, sadness, and surprise)[27,28]. Specifically, all 624 images in the stimulus set 1 were presented to the pretrained VGG-Face and their activations in the conv5-3 layer were extracted. First, we conducted a two-way non-repeated analysis of variance (ANOVA) with identity and expression as factors to detect units selective to facial expression (p ≤ 0.01) but not to face identity (p > 0.01). 1259 units out of the total 100,352 units (1.25%) in the conv5-3 layer were founded to have expression selectivity. Then, for each expression-selective unit, its tuning value [35] for each expression category was calculated to measure whether and to what extent it preferred specific expression. As shown in Fig. 1B, almost all units responded selectively to only one specific expression and exhibited a tuning effect. Finally, to test whether the responses of these expression-selective units provide sufficient information for successful expression recognition, we performed principal components analysis (PCA) on these unit activations to all images and selected the first six hundred principal components to perform an expression classification task, using a support vector classification (SVC) analysis with 104-fold cross-validation. The classification accuracy (mean ± standard error, 76.76% ± 1.59%) was much higher than the chance level (16.67%) and much higher than the accuracy with randomly shuffled expression labels (p = 1.8 × 10^−35^, Mann–Whitney U test)(Fig. 1C). The results indicated that the expression-selective units spontaneously emerged in the pretrained VGG-Face that was pretrained for face identity recognition, which echoed previous findings [19,20].

### 2. Human-like expression confusion effect of the expression-selective units in the pretrained VGG-Face

To examine the reliability of the expression-selective units, we used the classification model trained by stimulus set 1 to predict the expressions of images selected from the Radboud Faces Database (RaFD)[31]. The RaFD is an independent facial expression database including 67 face identities with different head and gaze directions, and only the front view images of each identity and each expression were used in the present study. The classification accuracy of the expressions from the RaFD was significantly higher than the chance level (accuracy = 67.91%, 95% confidential interval (CI) = [63.18%, 72.39%], image bootstrap with 10,000 replications)(Fig. 2A). It thus indicated that the expression-selective units in the pretrained VGG-Face had a reliable and general expression discriminability.

**Fig. 2.**
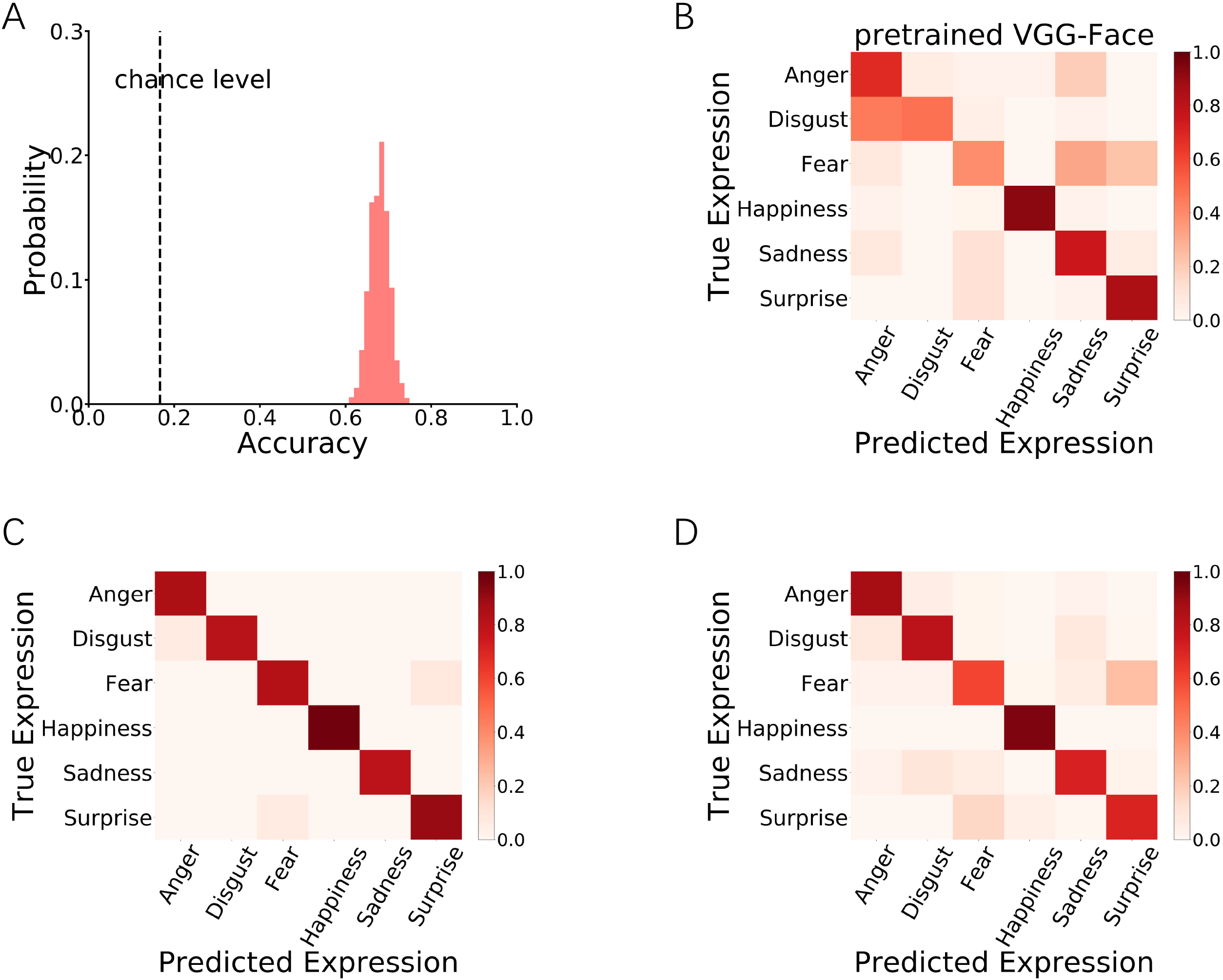
Human-like expression confusion effect of the expression-selective units in the pretrained VGG-Face. (A) The expression discriminability of the expression-selective units emerged in the pretrained VGG-Face for the RaFD. The black dashed line represents the chance level. (B) The confusion matrix of the expression-selective units for the RaFD. (C) Confusion matrix for the RaFD in human study (Adapted from Fig. 4 in Langner et al. 2010). (D) Confusion matrix for the NimStim set in human study (Adapted from Table 2 in Tottenham et al. 2009).

**Fig. 3.**
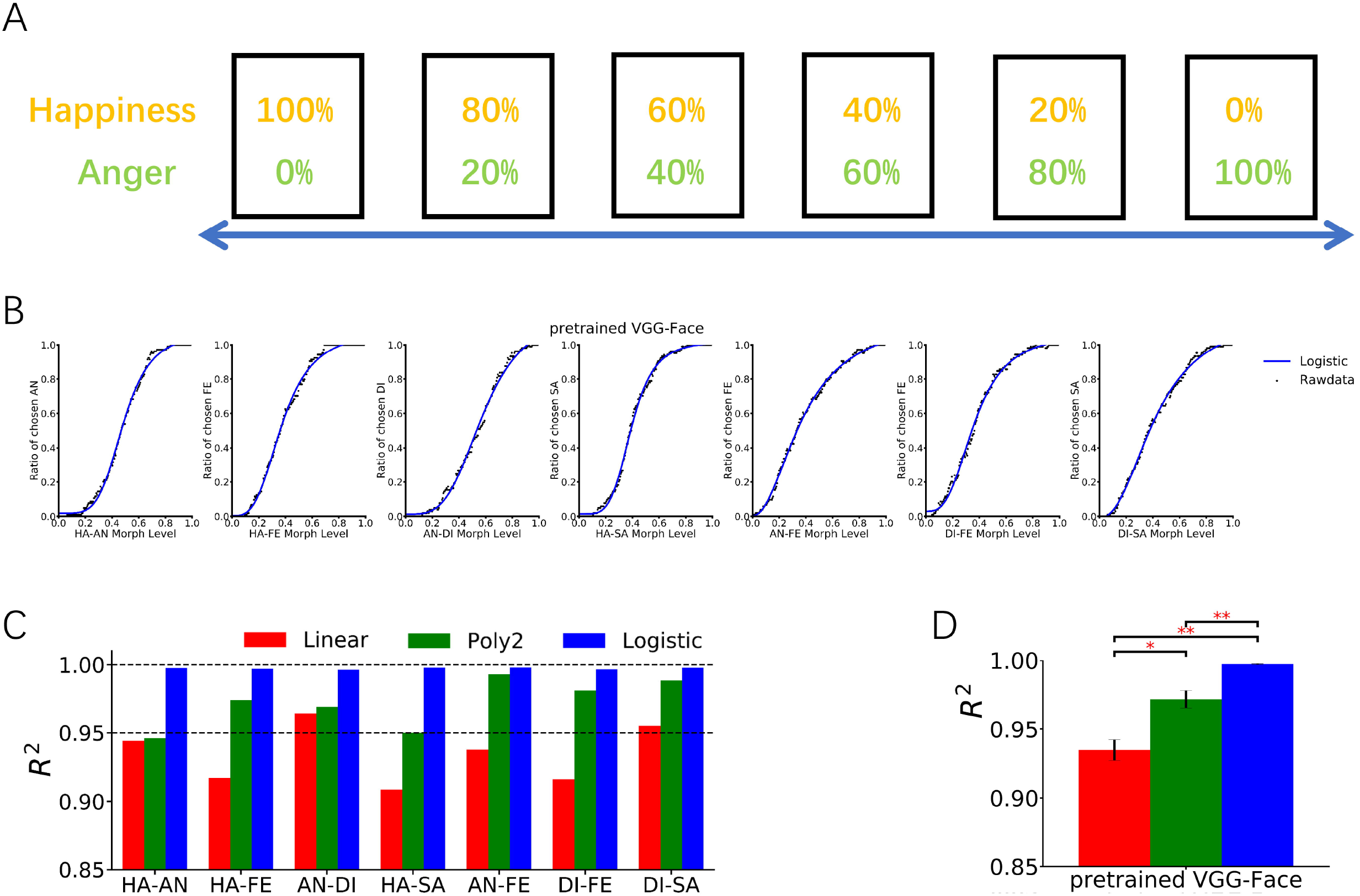
Categorical perception of facial expressions of the expression-selective units in the pretrained VGG-Face. (A) Example facial stimuli used in morph continua (Happy-Anger). (B) The identification rates for the seven continua. The identification rates refer to the identification frequency of one out of the two expressions. Labels along the x-axis indicate the percentage of this expression in facial stimuli. Black dots represent true identification rates. Blue solid lines indicate fitting for the logistic function. (C) Goodness-of-fit (R^2^) of each regression type for each expression continuum. The black dashed lines represent R^2^ at 0.95 and 1.00, separately. (D) Mean goodness-of-fit (R^2^) among expression continua. The R^2^ in the Logistic regression was much higher than the other two regressions. Error bars indicate SE. *: p≤0.05; **: p≤0.01; ***: p≤0.001.

Subsequently, to test whether the response profiles of these units were like the human behavior, we calculated a confusion matrix of the response patterns of these units to each expression of the RaFD (Fig.2B) and correlated it with the behavioral confusion matrix of facial expression recognition from one previous human study [31], which also used expression stimuli from the RaFD (Fig. 2C). We found that the two confusion matrices were highly correlated (r = 0.91, p = 1.6 × 10^−14^). The similarity in the confusing effects of facial expressions between VGG-Face and humans was further confirmed by using the behavioral confusion matrix from another human study [30], whose expression stimuli were selected from the NimStim database (r = 0.92, p = 4.1 × 10^−15^)(Fig. 2D). In all three confusion matrices, the true label of each expression category was the most frequently predicted label and consistent across matrices, and the confusion pattern in each matrix was almost coincided with each other. For example, like humans, fear and surprise may be confused with each other, and disgust was mistaken for anger more often than for other expressions. However, we found that these units also confused sadness with anger and fear, which was different from those in the human studies. Overall, the results suggested a similar expression confusion effect between the expression-selective units in the pretrained VGG-Face and humans.

### 3. Ecological validity of expression selectivity emerged in the pretrained VGG-Face

The facial expressions in the stimulus set 1 and RaFD were collected from the same actors and/or with detailed instructions to express different emotions, which thus had limited ecological validity. If the expression-selective units can recognize expressions, they should also be able to recognize the real-life facial expressions with ecological validity. To verify this, we performed the same analysis on expression images selected from the AffectNet database, which is a large real-world facial expression database [32]. We selected 4800 images with manually annotated expressions, 800 for each basic expression. Note that, in the AffectNet database, the face identities across expressions are distinct. Again, we found that these expression-selective units can classify facial expressions much better than chance level accuracy = 29.52%, 95% CI = [28.27%, 30.79%], image bootstrap with 10,000 replications)(Fig. 4A). Besides, their confusion matrix for facial expressions selected from the AffectNet database also showed significant similarities with those from humans (for RaFD, r = 0.62, p = 5.9 × 10^−5^; for NimStim, r = 0.67, p = 7.2 × 10^−6^)(Fig. 4B). The reliable human-like confusion effect of facial expression recognition suggested that the expression-selective units in the pretrained VGG-Face can recognize facial expressions in a way as humans do.

**Fig. 4.**
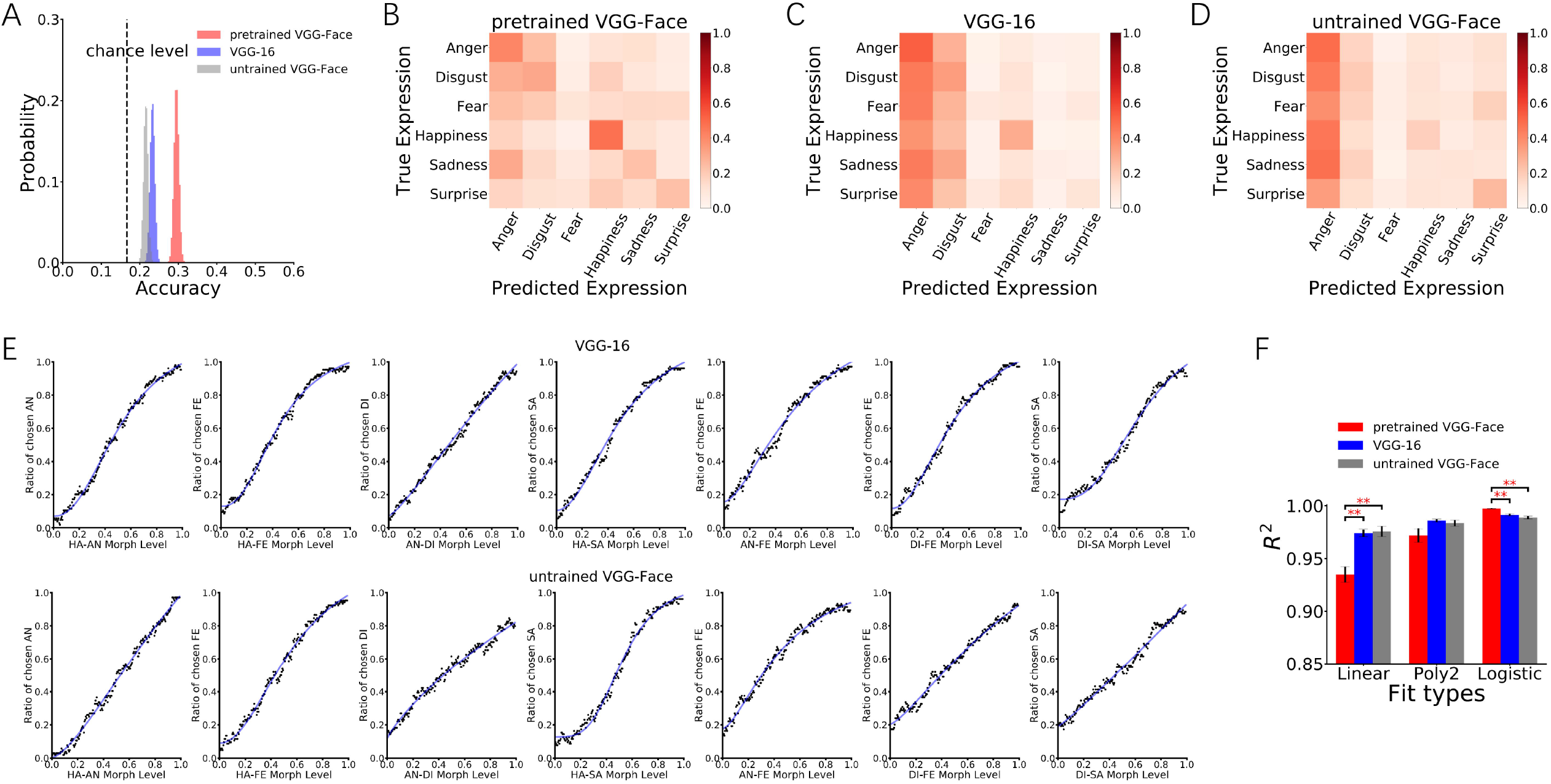
Expression recognition of the expression-selective units for the expressions from the AffectNet in the pretrained VGG-Face, VGG-16, and untrained VGG-Face. (A) The expression discriminability of the expression-slective units for each DCNN. Expression classification of the expression-selective units in the pretrained VGG-Face is much better than in the VGG-16 and untrained VGG-Face. The black dashed line represents the chance level. (B-D) The confusion matrix of the expression-selective units in the pretrained VGG-Face (B), VGG-16 (C), and untrained VGG-Face (D). (E) The identification rates for the seven continua in the VGG-16 and the untrained VGG-Face, respectively. Black dots represent true identification rates. Blue solid lines indicate fitting for the logistic function. (F) The Goodness-of-fit (R^2^) of each fit type for each DCNN. Logistic regression fits best for the pretrained VGG-Face than for the other two DCNNs, whereas linear regression fits worst for the pretrained VGG-Face. Error bars indicate SE. *: p≤0.05; **: p≤0.01; ***: p≤0.001.

### 4. The expression-selective units in the pretrained VGG-Face show human-like categorical perception for morphed facial expressions

One may argue that the similarity in the expression confusion effect does not necessarily mean expression-selective units perceive expressions in a human-like way. It might result from the similarities in physical properties of the expression images, since the image-based PCA analysis (i.e., principal components based on pixel intensities and shapes) could also yield a confusion matrix similar to that of humans [39]. Therefore, to further confirm whether these units could behave a human-like psychophysical response function to facial expressions, we tested whether their responses exhibited categorical effect of facial expression perception by using morphed expression continuum. Considering the generality of the categorical emotion perception in humans (Fig. 3A), we systematically tested the categorical effect in seven expression continua including happiness-anger, happiness-fear, anger-disgust, happiness-sadness, anger-fear, disgust-fear, and disgust-sadness. All of them have been tested in humans [36–38,42,40,41]. In detail, we designed a morphed expression discrimination task (Fig. 3B) that was comparable to the ABX discrimination task designed for humans [36–38]. The prototypic expressions were selected from the stimulus set 1. For each expression continuum, images of the two prototypic expressions were used to train an SVC model, and then the trained SVC model was applied to identify expressions of the morphed images. At each morph level of the continuum, the identification frequency of one out of the two expressions was defined as the units’ identification rate at the current morph level. We hypothesized that if the selective units perceive expressions like humans, i.e., showing categorical effect, then the identification curves should be S-shaped.

As predicted, for all continua, the identification curves of the selective units were S-shaped (Fig. 3C). To quantify this effect, we fitted Linear, Quadratic (Poly2) and Logistic functions to each identification curve, respectively. If the units exhibited a human-like categorical effect, the goodness of fit (R^2^) of the Logistic function to the curves should be the best. Otherwise, the goodness of fit of the Linear function to the curves should be the best if the units’ response followed the physical changes in images. As illustrated in Fig. 3D and 3E, we found that all seven identification curves showed typical S-like patterns (Logistic vs. Linear: p = 0.002; Logistic vs. Poly2: p = 0.002, Mann–Whitney U test).

### 5. The human-like expression perception only spontaneously emerges in the DCNN with domain-specific experience (pretrained VGG-Face), but not in those with domain-general visual experience (VGG-16) or without any visual experience (untrained VGG-Face)

So far, we had demonstrated that the human-like perception of expression could spontaneously emerge in the DCNN pretrained for face identity recognition. However, how did these express-selective units achieve human-like expression perception? Specifically, it was still unknown whether the spontaneous emergence of the human-like perception of expression depended on the domain-specific experience (e.g., face-related visual experience), a general natural object recognition experience, or even only the architecture of the DCNN.

To address this question, we introduced two additional DCNNs: VGG-16 and untrained VGG-Face. The architecture of VGG-16 is almost identical to the pretrained VGG-Face except that the last FC layer includes 1000 units rather than 2622 units. VGG-16 was trained for classifying 1000 object categories using natural object images from ImageNet [25], thus it only had object recognition experience. The untrained VGG-Face preserved the identical architecture of VGG-Face while randomly assigning the connective weights (Xavier normal initialization) [18,26], and had no training experience.

Images from the stimulus set 1 were also presented to VGG-16 and untrained VGG-Face respectively, and the responses of the units in the conv5-3 layer were extracted. Then, as that used in the pretrained VGG-Face, a two-way non-repeated ANOVA was performed to detect expression-selective units: 835 (0.83%) and 644 (0.64%) of the 100,352 total units were found to be expression-selective in the VGG-16 and in the untrained VGG-Face, respectively. It seemed that expression-selective units also spontaneously emerged in the VGG-16 with the experience of the natural visual objects, and even in the untrained VGG-Face without any visual experience.

Then, for each DCNN, images from the AffectNet Database were applied to test the reliability and generality of the units’ expression recognition ability. As in the pretrained VGG-Face, the classification accuracies in these two DCNNs were higher than the chance level (VGG-16: accuracy = 23.29%, 95% CI = [22.10%, 24.48%]; untrained VGG-Face: accuracy = 21.60%, 95% CI = [20.46%, 22.79%], bootstrap with 10,000 replications)(Fig. 4A). Crucially, by further comparing the classification accuracies of these two DCNNs with the pretrained VGG-Face, we found that the performance of the expression-selective units in the pretrained VGG-Face was significantly much higher than VGG-16 (p < 0.001, Mann–Whitney U test) and untrained VGG-Face (p < 0.001). Besides, the accuracy of expression-selective units in the VGG-16 was better than the untrained VGG-Face (p < 0.001). The results revealed expression-selective units in the DCNNs, whether with face identity recognition experience or not, could classify facial expressions. More importantly, the face identity recognition experience was more beneficial than general object classification experience for the enhancement of the units’ expression recognition ability.

Further and importantly, for both DCNNs, the similarities of expression confusion effect between the expression-selective units and humans were tested by correlating their confusion matrices with those of humans (Fig. 2C and 2D, respectively). The confusion matrix of expression-selective units in neither of the two DCNNs resembled those of humans (Fig. 4C, VGG-16: r_RaFD_ = 0.20, p_RaFD_ = 0.24; r_NimStim_ = 0.24, p_NimStim_ = 0.15)(Fig. 4D, untrained VGG-Face: r_RaFD_ = 0.18, p_RaFD_ = 0.29; r_NimStim_ = 0.20, p_NimStim_ = 0.23). Collectively, only the expression-selective units in the pretrained VGG-Face presented a human-like expression confusion effect. These results implied that, at least for the recognition facial expression, the domain-specific training experience was necessary for a DCNN to behave human-like perception.

As the expression-selective units in the VGG-16 and untrained VGG-Face showed no similarity to human expression recognition, we hypothesized that they may not perceive expressions like humans. To verify this hypothesis, we further investigated whether the expression-selective units in these two DCNNs would show categorical perception effect by used the same analysis mentioned above. As shown in Fig. 4E, the expression-selective units from both the VGG-16 and untrained VGG-Face only presented a weak S-shaped trend in very few continua. By comparing their goodness of fitting with that of the pretrained VGG-Face, the identification curves of the expression-selective units in the VGG-16 and untrained VGG-Face showed a more obvious linear trend than that in the pretrained VGG-Face (Linear: pretrained VGG-Face vs. VGG-16: p = 0.003; pretrained VGG-Face vs. untrained VGG-Face: p = 0.005; VGG-16 vs. untrained VGG-Face: p = 0.609, Mann–Whitney U test)(Fig. 4F). Correspondingly, they presented a worse logistic trend than the pretrained VGG-Face (Logistic: pretrained VGG-Face vs. VGG-16: p = 0.002; pretrained VGG-Face vs. untrained VGG-Face: p = 0.002; VGG-16 vs. untrained VGG-Face: p = 0.307)(Fig. 4F). Taken together, the face identity recognition experience, which was domain-specific, helped expression-selective units in the DCNN to achieve human-like categorical perception of facial expressions, whereas the general object classification experience and the architecture itself may only help capture physical features of facial expressions.

## Discussion and conclusion

The purpose of the current study was to evaluate whether the spontaneous emergence of human-like expression-selective units in DCNNs would depend on domain-specific visual experience. We found that the pretrained VGG-Face, a DCNN with irrelevant face identity recognition experience, could spontaneously generate expression-selective units. And these units allowed reliable human-like expression perception, including expression confusion effect and categorical perception effect. By further comparing the pretrained VGG-Face with VGG-16 and untrained VGG-Face, we found that although all the three DCNNs could generate expression-selective units, their performance of expression classification differed. The classification accuracy of the expression-selective units in the pretrained VGG-Face was the highest whereas that in the untrained VGG-Face was the lowest. More critically, only the expression-selective units in the pretrained VGG-Face showed apparent human-like expression confusion effect and categorical perception effect. Expression-selective units in both the VGG-16 and untrained VGG-Face did not perform similar to human perception, that is, they showed no human-like confusion effect and exhibited a continuous linear perception of morphed facial expressions instead of categorical perception. These results indicated that the human-like expression perception could only spontaneously emerge in the pretrained VGG-Face with domain-specific experience (i.e., visual experience of face identities), but not in the VGG-16 with task-irrelevant visual experience or the untrained VGG-Face without any visual experience. This finding supports the idea that human-like facial expression perception relied on face identity recognition experience.

Previous research has demonstrated that DCNNs could attain sensitivity to abstract natural features, such as number and face identity, by exposures to irrelevant natural visual stimuli [18,34,42], or even by randomly distributed weights without any training experience [21,43]. The DCNNs’ innate sense of number was consistent with the spontaneous representation of visual numerosity in various species, including nonhuman primates [44] and birds [45]. Similarly, the emergence of face identity selective units in untrained DCNNs was in line with the face selectivity founded in month monkey infants [46]. The spontaneous emergence of the selectivity of number and face identity in nonhuman and infant biological systems and untrained *in silico* DCNNs suggested that hard-wired connections of neural circuit were sufficient to perceive numerosity and face identity. Recent studies further explained the innate ability to recognize face identity might result from that face identity information could be represented by generic object features [18,47]. Therefore, in both biological systems and DCNNs, the extraction of high-order information of natural features such as number and face configuration depended much more on the physical architecture of the networks rather than to the training experience. Consistently, in the present study we found that besides the pretrained VGG-Face, both the VGG-16 and untrained VGG-Face could generate expression-selective units owing to the network architecture.

However, we argue that the pretrained VGG-Face is fundamentally different from VGG-16 and untrained VGG-Face. Unlike number sense and face identity recognition, only the information conveyed by the expression-selective units in the pretrained VGG-Face was sufficient to explain expression confusion effect and categorical expression perception observed in humans. Since the architectures of all three DCNNs (pretrained VGG-Face, VGG-16, and untrained VGG-Face) were identical, their divergence of expression perception originated from distinct training experiences. Namely, the human-like expression perception in the pretrained VGG-Face depended on domain-specific visual experience. The necessity of domain-specific training experience was consistent with the discoveries in biological systems. Specifically, Infants were not born with categorical perception of facial expressions. For instance, they began to show true categorical perception of happy faces and fearful faces only when they were at least 7-month old [48–50], and their discriminability of some other expression continua might develop even later [51]. In addition, a study examining international adopted children revealed that early postnatal deprivation to other-race faces disrupted expression recognition and heightened amygdala response to out-group emotional faces relative to in-group faces [52], revealing the importance of early domain-specific experience for the development of racial-specific facial expression processing. There is also evidence directly support that familiarity and perceptual learning can improve categorical perception [53,54]. Therefore, the architectures of both biological neural systems and artificial neural systems are insufficient to approach adult-level facial expression perception. Concurrent face identity development [55] or domain-specific training experience is needed.

Why would the human-like expression perception in DCNN distinctly rely on domain-specific experience compared to e.g., number sense and face identity recognition? We think that considering their distinct development or evolution in biological systems, the uniqueness of expression processing may originate from the difficulty of extracting abstract social information using generic natural features. The present findings revealed that while the expression-selective units in the pretrained VGG-Face extracted categorical/discontinuous expression information from morph continua, the expression-selective units in the VGG-16 and untrained VGG-Face merely extracted continuous linear information from visual features. The results indicated that the continuous representation of facial expression was architecture-dependent, but the categorical representation of facial expression was domain-specific experience dependent. Therefore, the face identity and facial expression were recognized by at least partly interconnected systems instead of separate parallel systems.

Indeed, the categorical perception of morphed expressions in the pretrained VGG-Face was in line with the categorical representation of expression in the amygdala, while the VGG-16 and untrained VGG-Face resembled pSTS which exhibited a continuous linear representation of morphed expressions [41,56]. The continuous representation of expression in the VGG-16 and untrained VGG-Face endowed the expression-selective units with a weak ability to recognize expressions, coinciding with the previous finding showing recognition of expressions at a certain level due to image-based principal components [39]. Thus, it indicated that expression representation in the ventral visual pathway, VGG-16 and untrained VGG-Face relied mainly on the similarities of physical properties of images with facial expressions, whereas expression representation in the amygdala and pretrained VGG-Face depended on the categorical information in faces (namely, social meaning) that was critical for making rapid and correct physiological responses to threat and danger. Based on these findings, we would suspect that the function of the core face network may be inborn, but the normal function of the extended face network would rely on postnatal domain-specific experience. However, future developmental study combined with advanced neuroimaging techniques for infants may be needed to confirm this hunch.

Together, theoretical contributions of the present study are two-folds. Firstly, our findings added strong evidence supporting DCNNs’ potential to perform human-like representation. The spontaneous generation of human-like facial expression confusion effect and categorical perception in the pretrained VGG-Face were in line with other similarities between DCNNs and humans, such as similar coding hierarchy as the feedforward visual-cortical network [16,17,57,58], number sense [34,43], object shape perception [59], face identity recognition [18,21], and perceptual learning [60]. Secondly and perhaps more importantly, our computational findings revealed the necessity of domain-specific visual experience of face identity for the development of facial expression perception, and suggested a biologically plausible model for internal brain processing of social information in addition to generic natural features after pretrained on domain-specific tasks, highlighting the contribution of nurture to form human-like facial expression perception. As there exists challenges to conduct human developmental research, such as ethical concerns, recruitments difficulties, participant attrition, and time-consuming (particularly in long-term longitudinal studies), the advantage of systematic comparisons among DCNNs at different levels showcases how DCNN could be appropriately utilized as a powerful tool for the study of human cognitive development. Beyond the weak equivalence between human and DCNNS at the input-output behavior, emerging simulated algorithms between models and humans could be established through domain-specific experience.

## Acknowledgements

This work was supported by the National Key R&D Program of China (2019YFA0709503), National Nature Science Foundation of China grant (31671133), National Basic Research Program.

## Author contributions

L. Zhou and K. Zhou contributed to the conceptualization and methodology. L. Zhou contributed to the data curation and writing-original draft. K. Zhou and M. Meng contributed to the writing-review & editing and supervision of the project.

